# Evaluation of mouse behavioral responses to nutritive versus nonnutritive sugar using a deep learning-based 3D real-time pose estimation system

**DOI:** 10.1101/2022.09.19.508605

**Authors:** Jineun Kim, Dae-gun Kim, Wongyo Jung, Greg S. B. Suh

**Affiliations:** Department of Biological Sciences, Korea Advanced Institute of Science and Technology (KAIST), Daejeon 34141, Republic of Korea

## Abstract

Animals are able to detect the nutritional content of sugar independently of taste. When given a choice between nutritive sugar and nonnutritive sugar, animals develop a preference for nutritive sugar over nonnutritive sugar during a period of food deprivation^1-5^. To quantify behavioral features during an episode of licking nutritive versus nonnutritive sugar, we implemented a multi-vision, deep learning-based 3D pose estimation system, termed the AI Vision Analysis for Three-dimensional Action in Real-Time (AVATAR)^6^. Using this method, we found that mice exhibit significantly different approach behavioral responses toward nutritive sugar versus nonnutritive sugar even before licking a sugar solution. Notably, the behavioral sequences during approach toward nutritive versus nonnutritive sugar became significantly different over time. These results suggest that the nutritional value of sugar not only promotes its consumption, but also elicits distinct repertoires of feeding behavior in deprived mice.

## Introduction

Animals innately prefer nutritive sugar over nonnutritive sugar or can learn to develop this preference over time^1,3,7^. Given that sweet-blind mice prefer nutritive over nonnutritive sugars, this preference is independent of taste input^8-10^. It is unclear, however, whether an increase in the consumption of a nutritive sugar over a nonnutritive one is a behavioral response that can be quantified and analyzed in detail. Mice may superficially appear to approach nutritive sugars differently than they approach nonnutritive sugars, but the details of their behavioral repertoires spurred by metabolic needs may be different. Indeed, superficially similar behaviors could be distinguished by analyzing the behavior itself using precise behavioral quantification methods^11,12^.

Innately motivated behaviors, including the consumption of nutritive sugars during a period of food deprivation, are classically composed of three main phases: appetitive, consummatory, and satiety phases^13,14^. It is well known that animals display significantly different sequences of behavior during each phase, each often requiring a massive amount of behavioral annotation. Recent advances in computer vision machine learning have led to an efficient and precise way to extract the postures of a behaving animal from each image at a high spatial and temporal resolution^15-17^. In studies of the motor system, for example, a deep learning tool with a higher resolution has been used to analyze the role of the motor cortex^18,19^. Recently, the pose estimation model was used to analyze complex social behaviors of mice that include mating and fighting^11,17^. The mounting behavior of a male mouse, for example, varies with the sex of the encountered mouse. A male mouse may mount both male and female mice, but the sequence of his mounting behavior is distinct enough for a machine-learning-based classifier to distinguish between mounting followed by attack versus mating^11^. It is unknown, however, whether mice exhibit distinct appetitive-related behaviors in response to the nutritional value of food.

In this study, we applied a deep learning-based 3D pose estimation system, the AI Vision Analysis for Three-dimensional Action in Real-Time (AVATAR) system to use the coordination of 9 body points to quantify and analyze animal behaviors^6^ while they develop a preference for nutritive sucrose over nonnutritive sucralose. We found that fasted mice not only consumed considerable amounts of sucrose solution, but exhibited significantly different approach behavioral repertoires toward sucrose solution within 30 minutes of the first exposure. Using a deep learning-based system combined with a conventional measurement of food consumption, we were able to shed light on how nutritive food is selected over equally palatable, yet nonnutritive food by fasted mice and create a precise and large-scale behavioral dataset.

## Results

### Mice develop a preference for nutritive sugar over nonnutritive sugar when fasted

Previous studies using *Drosophila* revealed that flies develop a preference for nutritive sugar depending on their energy status. Fasted flies select nutritive sugars over non-nutritive sugars within 5 minutes, whereas sated flies do not necessarily demonstrate a preference for nutritive sugar ^4,5^. It is unknown, however, whether mice that had not been previously exposed to nutritive or nonnutritive sugars exhibit the starvation-induced preference for nutritive sugars. To investigate this matter, we first established a behavioral assay to measure the starvation-induced preference for nutritive sugars. To determine whether mice can detect and prioritize the nutritional content of sugar over its sweet content without conditioning, we designed a two-choice assay in which naïve mice (i.e., mice that had not been exposed to a plain sucrose or sucralose solution) were given a choice between a bottle containing a 100-mM sucrose solution and a bottle containing an equally sweet 0.5-mM nonnutritive sucralose solution under fasted conditions.

Overnight fasted mice were presented with the two bottles and their consumption was measured using a custom-made lickometer. Within 30 minutes, fasted mice consumed more of the sucrose solution than the sucralose solution (**Figure 1B**). We also observed that fasted mice rapidly developed a preference for sucrose within 10 minutes (**Figure 1B, black arrow**). The inter-bout interval was shorter while licking the sucrose solution compared to the sucralose solution (**Figure 1E**). These results suggest that fasted and unconditioned mice rapidly develop a preference for sucrose over sucralose.

**Figure 1.**
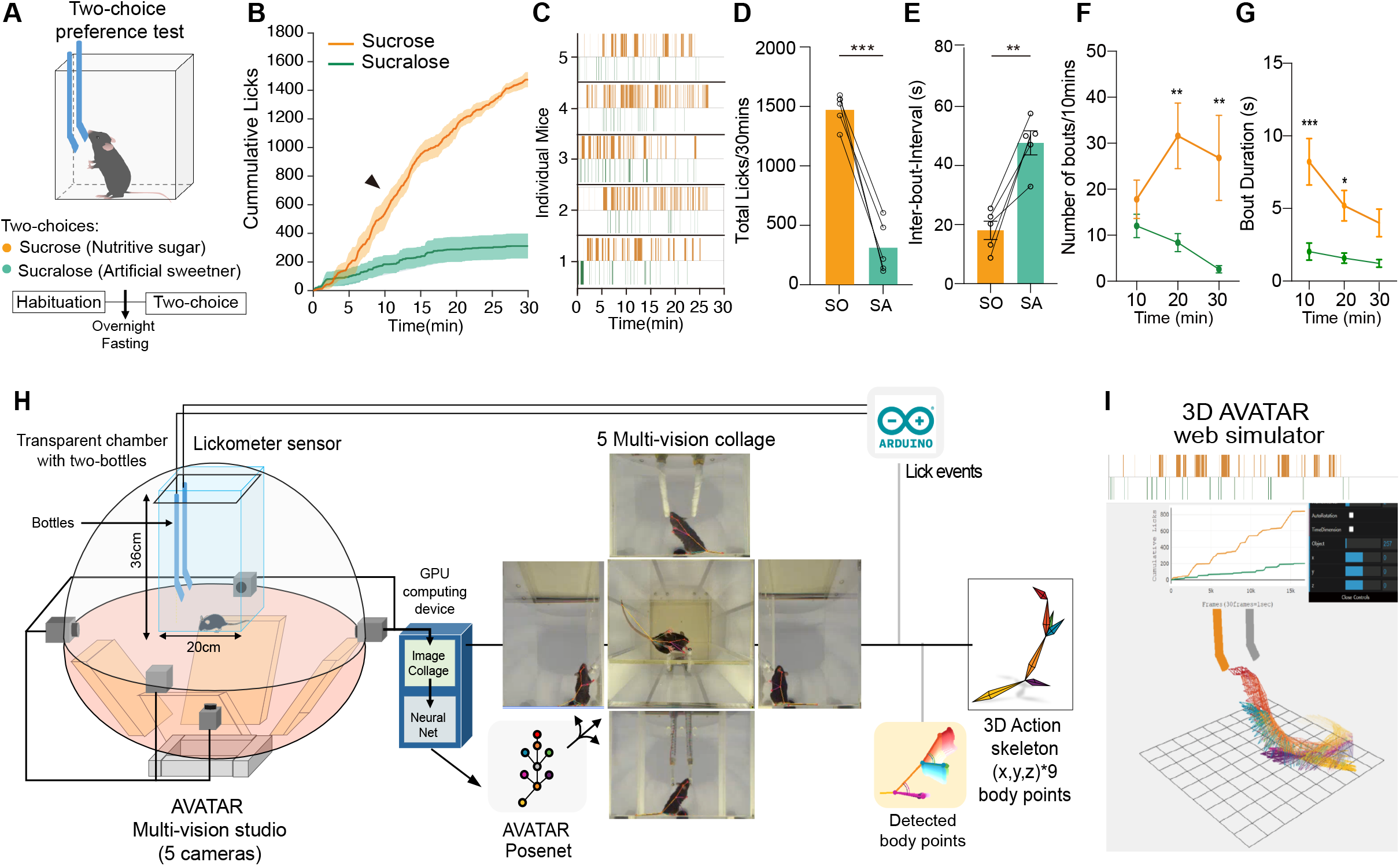
Mice rapidly develop a preference for nutritive sugar when fasted. **(A)** Left: Schematics of two-bottle choice preference for nutritive sucrose (100mM sucrose) and non-nutritive sucralose (0.5mM sucralose) in mice fasted overnight. **(B)** Plots for cumulative licks over 30 mins of two-choice preference assay. Orange line: sucrose, Green line: sucralose. Lines and shaded areas indicate mean +/- SEM. **(C)** Raster plots for lick event train for each analyzed mouse. Orange ticks: licks for sucrose, Green ticks: licks for sucralose. (**D-G**) Quantifications of total licks (D), interbout-interval (E), number of lick bouts, and duration of lick bout in 10-minute time bin during 30 mins of the assay (F and G). Lick bout was defined as a series of licks with an inter-lick interval of less than 1 second. **(H)** Schematics of a two-bottle choice paradigm in the AVATAR 3D studio. Two bottles are connected to a custom-made capacitance-based lickometer. Licking events and 5 multi-vision collage images were synchronized and 27 arrays of the 3D action skeleton were collected. **(I)** Screenshot of the web simulator that can visualize the dynamic 3D pose sequence and lickometer sensor data while using AVATAR. A link to the AVATAR web simulator during the two-choice preference task: <u>Two choice.js app</u>.

### Fasted mice exhibit significantly different behaviors toward nutritive sucrose versus nonnutritive sucralose

We next used a machine-learning-based method to quantify and classify changes in the sequence of behaviors as they developed a preference for sucrose. Our analysis of licking behavioral patterns revealed that the number of licking bouts for sucrose increased rapidly while the duration of licking bouts decreased (**Figures 1F and 1G**), suggesting an increase in preference for sucrose during the appetitive phase^13,20^. We hypothesized that the behavioral changes during the appetitive phase may coincide with the development of sucrose preference.

We therefore sought to track and record distinctive features of the appetitive behavior toward sucrose versus sucralose by using a deep learning-based 3D pose-estimation algorithm (AVATAR)^6^. To do so, we constructed a system to measure and quantify behavior sequences during the two-choice assay. We first devised the two-bottle choice task chamber and lickometer, in which two licking spouts on the bottle are exposed on a glass wall in the chamber. The behaviors of mice were recorded from five different directions in the AVTAR multi-vision studio (**Figure 1H**). Licking events and 5 multi-vision collage images were temporally synchronized. We then collected the coordination of body pose (27 arrays; 9 body-points x 3 axes) at each video frame using the AVATAR posenet that was trained for approximately 20 frames with the two-bottles (**Figure 1H**). We next generated a 3D action skeleton, which is illustrated in the AVATAR web stimulator (**Figure 1I**), to visualize and analyze the dynamic action sequences and cumulative licking events.

To analyze the behavioral sequences selectively during the phase of approach behavior, we collected the coordination of body pose for approximately 3.3 seconds (100 frames with 30 FPS; 2700 vectors x the number of licks) before each licking bout. Several binary classification algorithms (Naive Bayes, SVM with Gaussian Kernel, and logistic regression classifiers)^21^ were used to evaluate the behavior sequence data and allow us to discriminate between the approach behaviors for sucrose versus sucralose (**Figure 2B**). We found that the performance of the Naive Bayes and Gaussian SVM classifiers was significantly higher than by chance (Naive Bayes: 75.7%, SVM Gaussian: 81.2%) (**Figures 2B-2D**). Remarkably, both classifiers showed higher classification accuracy when the behavior sequences extracted from the late period of the same experiment were used (**Figure 2E**). This suggested that the approach behavior toward sucrose is qualitatively distinct from the behavior toward sucralose over time.

**Figure 2.**
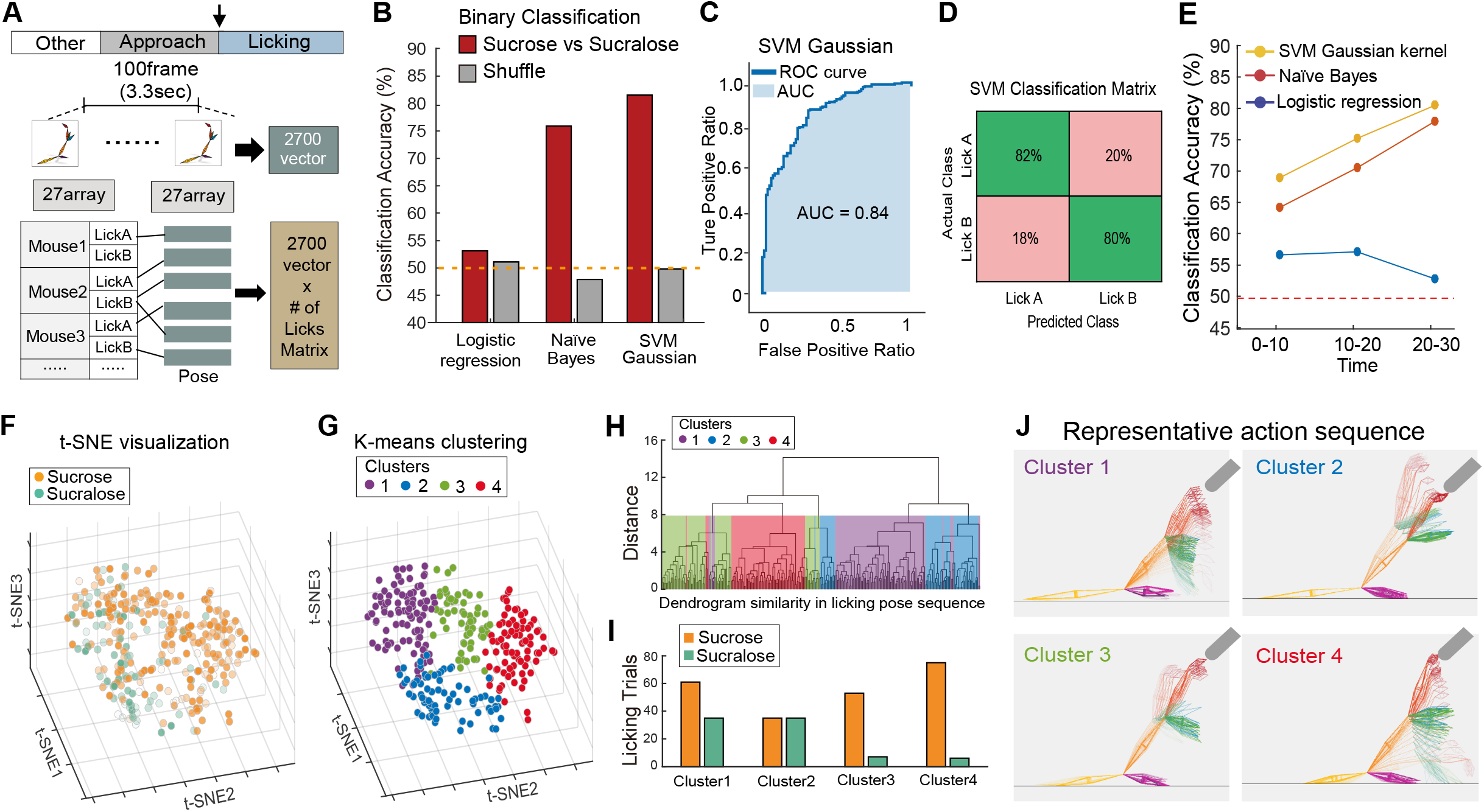
Analysis of two-bottle choice behaviors using the AVATAR system. **(A)** For the pose sequence data aligned to the licking events, we gained 27 arrays (x,y,z points from 9 body points) at each frame for 100 frames before the onset of licking bouts. The action skeleton array (pose sequence data) during the target time window, as one-hot vector of 2700, was used for each mouse as the final matrix data to be further analyzed. **(B)** Bar graphs indicating the classification accuracy by binary classifiers with shuffled (grey) and labeled data (red); Logistic regression (Label:53.1%, Shuffle:51.1%), Naïve Bayes (Label:75.7%, Shuffle:47.8%), SVM with Gaussian kernel filter (Label:81.2%, Shuffle:49.8%). The orange dash line denotes 50% accuracy as a reference. **(C)** The receiver operator characteristic (ROC) curve from the SVM classifier in **B**. The Area Under the Curve (AUC) for the ROC curve is 0.84. **(D)** The confusion matrix of the SVM classification is used to evaluate the performance of the classification model for the predicted and actual classes. **(E)** Classification accuracy in 10-minute intervals (3 phase: 0-10 min, 10-20 min, 20-30 min). Yellow line: SVM Gaussian kernel, Redline: Naive Bayes, and Blue line: Logistic regression; 0-10(min) phase: Yellow=69.4, Red=64.5, Blue=57.0; 10-20(min) phase: Yellow=75.6, Red=70.8, Blue=57.3; 20-30(min) phase: Yellow=81.2, Red=78.3, Blue=53.4. **(F)** Three-dimensional projection representing clusters by visualizing t-SNE for pre-lick pose sequence data, labeled with each licking bottle. Orange dots: licking from sucrose. Green dots: licking from sucralose. The scale of each color saturation implies a time flag during the task (Saturation: 10∼100% = Frame: 1∼60000). Total cluster points consisted of 307 licking trials (sucrose: 224 points, sucralose: 83 points). **(G)** Three-dimensional projection representing each cluster classified by the unsupervised algorithm (K-means classification). Cluster 1: Purple, Cluster 2: Blue, Cluster 3: Green, Cluster 4: Red. **(H)** Dendrogram chart to illustrate the similarity and hierarchical relationship of all 307 pose sequences. They are divided into 4 sub-clusters with 10.5 similarities and compared with the t-SNE cluster in **G** through the color box tag. Hierarchically, clusters 1 and 2 are close, and clusters 3 and 4 are close. **(I)** Bar graphs demonstrating the licking trials of each t-SNE cluster. Cluster1 has 61(63%) sucrose trials and 35(47%) sucralose trials; Cluster2: 35(50%) sucrose, 35(50%) sucralose; Cluster3: 53(88%) sucrose, 7(12%) sucralose; Cluster 4: 75(93%) sucrose, 6(7%) sucralose. **(J)** Representative 3D action sequence patterns from each cluster identified in **G**. Top left (Cluster #1): heads up toward the sipper, Top right (Cluster #2): heads down, Bottom left (Cluster #3): stretching its neck toward the sipper, Bottom right (Cluster #4): running toward a sipper. A <u>link to the AVATAR simulator for each cluster</u>.

To determine how the approach behavior toward sucrose versus sucralose changed over a period of 30 minutes, we embedded the detected body postures during approach to each sugar into a 3D space by using a t-distributed stochastic neighbor (t-SNE) for visualization. In the 3 dimensional t-SNE space, we found that the body postures during the approach behavior were distributed separately (**Figure 2F**). Furthermore, we observed that the difference in behavioral response between the two approach behaviors substantially increased over time (**Figure 2F**). This is consistent with the improvement of binary classifier’s performance trained with the approach behavior during the late period of the two-choice assay (**Figure 2E**). Unsupervised k-means clustering on the t-SNE embedding further revealed that the approach behavior was clustered into 4 types (**Figure 2G**). Clusters 3 and 4 mainly represented the approach behavior specific for sucrose (**Figure 2I**). To identify any similarities among the 4 clusters, we applied the hierarchical clustering method, which consisted of an unweighted pair group method with arithmetic mean (UPGMA), and plotted a dendrogram (**Figure 2H**). Notably, Clusters 3 and 4 had the greatest similarities in the approach behavior during the late period of the two-choice assay (**Figures 2F-2H**). These results indicated that an approach behavior toward nutritive sugar was significantly different from another approach behavior toward nonnutritive sugar.

We further explored the features of the sucrose-specific approach behavior. To display representative behavioral sequences in each cluster, we reconstructed the representative approach behavior of each cluster using a centroid of the k-means clustering into a 3D posture (**Figure 2J**). Interestingly, the representative behavior of Cluster 3, which largely comprised the approach behavior toward sucrose, consisted of initially selecting the opposite bottle and then switching to the sucrose bottle (**Figure 2J left bottom)**. The representative behavior of Cluster 4, another sucrose-approach cluster, consisted of the mice directly raising their head to the sucrose bottle from the bottom of the chamber, rather than approaching from the top (**Figure 2J right bottom**). Inferred from the qualitative differences in the approach behavior, segregated neural circuits may be activated in order to trigger distinct feeding behaviors toward sucrose versus sucralose, as suggested in the previous studies^2,3^.

## Discussion

In our attempt to identify differences in the approach behavior of fasted mice toward nutritive sugar versus nonnutritive sugar, we found that fasted mice can rapidly recognize and select nutritive sucrose over nonnutritive sucralose without conditioning. Using the deep-learning 3D pose estimation model and the machine learning based-classifiers, we identified a significant difference during the approach behavior toward sucrose versus sucralose. Notably, the difference developed within 30 minutes of exposure. Furthermore, among two distinct types of the approach behavior revealed by unsupervised clustering, the sucrose-specific approach behavior became prevalent over time.

Researchers have traditionally measured the amounts of consumed food or water to determine feeding or drinking behaviors^22,23^. In this study, we combined these conventional measurements with the quantification of mice behaviors using machine-learning approaches, including 3D pose estimation, supervised learning algorithms for classifications, and dimensionality reduction. These methods allowed us to evaluate the behavior of mice during the appetitive phase with precision and revealed that fasted mice exhibit qualitatively different approach behavior according to their metabolic needs. It would be interesting further to elucidate how two distinct approach responses for sucrose and sucralose are controlled by different neural circuits.

Recent advances in the deep learning-based pose estimation have introduced a new way of analyzing detailed behavioral responses^24-30^. Instead of providing a subjective analysis, it can automatically quantify animal behaviors at a high resolution to produce a large and highly accurate dataset^11,30^. In this study, we used the recently developed AVATAR, which facilitated the creation of an accurate 3D pose estimation of mice during the two-choice assay^6^. This allowed us to overcome the limitation of the 2D pose estimation model with a single camera view. The multiple directions of recording used in our method permitted the detection of each body posture with precision despite potential occlusion by the spout of bottles.

For future studies, we plan to use this method to quantify and analyze other goal-directed behaviors, including eating, lever pressing, or nose poking, to further understand the dynamics of these behaviors and gain insight into neural substrates that are tuned to these behaviors. Furthermore, the two-bottle choice paradigm adopted in the AVATAR studio would be applicable for providing quantifiable behavioral scoring of affective behaviors caused by neurological disorders – depression, anhedonia and post-traumatic stress disorder (PTSD) in rodent models.

## Methods

### Animals

For wild-type experiments, male C57BL/6J (Jackson Laboratories stock #000664) adult mice (8-16 weeks of age) were used. All mice were single housed under a 12 h light/dark cycle and *ad libitum* access to food and water All animal experiments were performed according to protocols approved by KAIST IACUC following the National Institutes of Health guidelines for the Care and Use of Laboratory Animals.

### Short-term two-bottle preference test

For three days, single-housed naive mice were acclimated to two bottles with drinking spouts in their home cage. Mice were habituated in a behavior apparatus (20 × 20 × 20 cm) with two spouts connected to a custom-designed lickometer (Arduino UNO) before testing. Mice were trained to lick from bottles with spouts by restricting their access to water for 20 hours and then introducing them to the cage for two days of habituation. With no side bias for each sipper - less than a 25% preference index - and at least 200 licks within thirty minutes, these acclimation sessions were considered successful. They are then placed in a home cage for one day and fed *ad libitum* with water and food. Mice were housed in a new bedding cage and deprived of food for 18-20 hours with free access to water for fasting conditions. Mice were acclimated in the behavior chamber box for 20 minutes before being placed into an apparatus with two bottles containing either 100 mM sucrose or 0.5 mM sucralose. All behavioral experiments were videotaped and recorded 1-2 hours after the dark cycle started. Preference index (%); percentage of the total licks for sucrose or sucralose = (Lick_Sucrose_ – Lick_Sucralose_)/(Lick_Sucrose_ + Lick_Sucralose_) × 100. The raw data from the lickometer was used to investigate licking dynamics using custom-designed MATLAB code.

### AVATAR 3D system

A specially designed multi-vision system was used to quantify the mice’s 3D behavior. The vision system, which consists of four cameras on the sides and one on the bottom, can closely observe the mouse’s external shape. Each camera’s image data is sent to a PC for analysis and concatenation into a single image frame. AVATAR posenet analyzes these frames (3600×2000 pixel), detecting 9 body-points (nose, head, body center, anus, forelimbs, hind limbs, tail tip) of the target mouse in each area. Through the 3D reconstruction algorithm, these 2D coordinates are calculated as XYZ coordinates where the mouse was actually located. The final output data is a csv file containing a recorded frame (54000 raws; 30min) and body coordination information (27 arrays; 9 body points x 3 axes). To synchronize the behavior recording with the custom-made lickometer, a custom-made lickometer sent the specific signal on the camera’s image whenever the mice licked the bottle, and the AVATAR posenet further detected the specific signal from the camera’s image.

### AVATAR Web Simulator

The AVATAR web simulator can visualize the pose data file (csv) quantified in the AVTAR 3D system. The 9 body-points coordination are combined into 8 vector sets that represent the actual mouse skeleton (nose-head, head-body center, body center-anus, anus-tail tip, body center-left forelimb, body center-right forelimb, anus-left hindlimb, anus-right hindlimb). The webGL-based 3D visualization library is used to simulate these vector data sets. This web simulator allows researchers to observe the skeletal changes and behavior of mice. It is possible to control the time frame and pose sequence time-bin, view-point, and pose records. It also visualizes the actual bottle position and dynamically plots the lick data of each bottle consumed by the mouse using lickometer sensor data.

### Analysis

For the pose sequence classification, the “Classification Learner App” from the Statistics and Machine Learning Toolbox for MATLAB R2022a was used. We selected several classifiers, such as Naive Bayes, SVM with Gaussian Kernel, and logistic regression. The software allowed users to explore data, choose features, and configure validation strategies using the Classification Learner App. Confusion Matrix plots, Parallel Coordinates plots, ROC Curve plots, Scatter plots, and the classification accuracy of the model developed are all generated using this interactive classifier. Also, t-SNE clustering, k-means classification, and hierarchical clustering method (UPGMA) analysis results were solved using MATLAB.

### Quantification and Statistical Analysis

All statistical analysis is done in MATLAB or Prism software. An unpaired or pair-wise comparison was made using a two-tailed student’s t-test. Data are presented as mean ± s.e.m. All statistical analyses were performed with GraphPad Prism 9.0.2. To compare two groups that present the normal distribution, unpaired two-tailed t-tests were conducted.

## Data and Software Availability

The datasets that support the findings of this study are available from the corresponding author upon reasonable request.

## Acknowledgements

This study was supported by grants from KAIST Advanced Institute-X (KAIX) Fellowship and ASAN Biomedical Science Fellowship to J.K, grants from the Samsung Science and Technology Foundation (SSTF-BA-1802-11), National Research Foundation of Korea (NRF-2020R1A2C2009865 and NRF-2022M3A9F3082982) to G.S.B.S. and (NRF-2019M3E5D2A01066259) to Daesoo Kim who contributed to the development of the AVATAR system, the AI-based analysis and interpretation of animal behavior, and a grant from the NRF funded by Ministry and Science and ICT (2021 NRF-M3F3A2A01037365) to G.S.B.S as a co-PI.

## Author Contributions

D-G. K. developed the AVATAR hardware and software algorithm platforms. J. K. performed behavioral experiments with help from W.G. and D-G. K. J.K. and W.G. contributed to the development of the hardware and analysis platform for customized lickometer. G.S.B.S conceived and supervised the project. J.K., and G.S.B.S wrote the manuscript with inputs from other authors.

